# Worth the work? Monkeys discount rewards by a subjective adapting effort cost

**DOI:** 10.1101/2023.01.10.523384

**Authors:** Mark Burrell, Alexandre Pastor-Bernier, Wolfram Schultz

**Author notes:** **Corresponding Author:** Wolfram Schultz. **Author Contributions:** MB, AP-B and WS designed the experiment, MB conducted experiments and analyzed data, MB wrote the paper. **Competing Interest Statement:** The authors declare no competing interests.

## Abstract

All life must solve how to allocate limited energy resources to maximise benefits from scarce opportunities. Economic theory posits decision makers optimise choice by maximising the subjective benefit (utility) of reward minus the subjective cost (disutility) of the required effort. While successful in many settings, this model does not fully account for how experience can alter reward-effort trade-offs. Here we test how well the subtractive model of effort disutility explains the behavior of two non-human primates (*Macaca mulatta*) in a binary choice task in which reward quantity and physical effort to obtain were varied.Applying random utility modelling to independently estimate reward utility and effort disutility, we show the subtractive effort model better explains out-of-sample choice behavior when compared to parabolic and exponential effort discounting. Furthermore, we demonstrate that effort disutility is dependent on previous experience of effort: in analogy to work from behavioral labour economics, we develop a model of reference-dependent effort disutility to explain the increased willingness to expend effort following previous experience of effortful options in a session. The result of this analysis suggests that monkeys discount reward by an effort cost that is measured relative to an expected effort learned from previous trials. When this subjective cost of effort, a function of context and experience, is accounted for, trial-by-trial choice behavior can be explained by the subtractive cost model of effort.Therefore, in searching for net utility signals that may underpin effort-based decision-making in the brain, careful measurement of subjective effort costs is an essential first step.

**Significance:** All decision-makers need to consider how much effort they need to expend when evaluating potential options. Economic theories suggest that the optimal way to choose is by cost-benefit analysis of reward against effort. To be able to do this efficiently over many decision contexts, this needs to be done flexibly, with appropriate adaptation to context and experience. Therefore, in aiming to understand how this might be achieved in the brain, it is important to first carefully measure the subjective cost of effort. Here we show monkeys make reward-effort cost-benefit decisions, subtracting the subjective cost of effort from the subjective value of rewards. Moreover, the subjective cost of effort is dependent on the monkeys’ experience of effort in previous trials.

## Introduction

Reward follows work, but what reward is worth the work? This subjective decision is fundamental to understanding animal and human behaviors during effort-based decision-making. Notably effort requirements can vary wildly, whether it be walking to the kitchen for breakfast, running for a train to work or hiking up a mountain on holiday. The brain must be able to compare all the effort costs in choosing these actions against their potential rewards and therefore should possess flexible mechanisms for effort-based decision-making.

When consumer choice theory and labour supply theories are used to understand consumer and worker behavior, it is assumed decision makers balance the possible utility (subjective benefit) against the disutility (subjective cost) of effort, with consumer choice theory predicting choices are made simply on net utility (reward minus cost, Fehr and Goette, 2007). Experimental results from optimal foraging theory, psychology and economics all support this conclusion (Bautista et al., 2001; Fehr and Goette, 2007; Zipf, 2016). However, in several contexts this model fails to explain why humans and animals select more effortful options (Friedman et al., 1968; Clement et al., 2000; Lydall et al., 2010). In animals, both context and order of experience serve as references that significantly affect reward-effort trade-offs (Knauss et al., 2020).

An attractive explanation of these phenomena is the reference dependency of effort-based decision-making developed in human behavioral labour economics (Fehr and Goette, 2007; Abeler et al., 2011; Crawford and Meng, 2011): choices are not made on the absolute effort costs but relative to a conditional expectation of effort. In this paradigm, there exists an internal expectation of effort (‘effort reference-point’) informed by context and experience. Increases in this reference point increase willingness to pick effortful options as the relative magnitude of the reference-dependent effort cost decreases. Furthermore, responses may differ dramatically on either side of the reference point: the work-decisions of New York taxicab drivers were found to be six-fold more sensitive to above-expected workloads as to below-expected workloads (Crawford and Meng, 2011). Similar findings of asymmetric work responses have been made in the economic literature (e.g.Akerlof and Yellen, 1990) and in animals (e.g. van Wolkenten et al., 2007).

However, little economic work has been done to study how effort reference-points change over the course of time in a single task, rather focusing on identifying what forms an appropriate reference point or modelling specific situations using reference-dependent preferences (De Giorgi and Post, 2011; Allen et al., 2016). Within prospect theory, a related theory of reference-dependent reward preferences, a few studies have documented changes in reference points in single task contexts in response to wins/loss history, supporting a role for experience in determining reference points (Köszegi and Rabin, 2006; Arkes et al., 2008).

Here we were interested in measuring effort disutility in rhesus macaques as a foundation to establishing the neurobiological basis of the reward-effort trade-offs. Previous tasks used to measure effort-costs have often additional confounds that make it difficult to directly assess the subjective value of effort. For example, in both commonly used designs in rodent studies, lever-pressing and the T-maze, high effort options are correlated with longer delays from trial onset (Salamone et al., 1994; Cousins et al., 1996). Given the extensive work demonstrating the subjective cost of time (temporal discounting) in both behavioral experiments and encoding by various brain regions (Critchfield and Kollins, 2001; Kobayashi and Schultz, 2008; Prévost et al., 2010) such designs are not suitable for isolating the subjective cost of effort from time. We apply a random utility model to independently model reward utility and effort disutility from safe options (Mcfadden, 1973; McFadden and Train, 2000). From the estimates of reward utility and effort disutility we demonstrated through out-of-sample testing that the monkeys’ choices closely follow those of a utility maximiser employing a subtractive model of effort costs. However, the willingness to exert effort was greater in both animals after effort expenditure, which can be explained by a model in which the reference point is learnt based on recent experience.

## Materials and Methods

The experiments were conducted on two purpose-bred male rhesus macaques (*Macaca mulatta*): Monkey U (weighing 14.5 kg) and Monkey W (weighing 11.5 kg). This research (PPL 70/8295) was ethically reviewed, approved, regulated and supervised by the following United Kingdom (UK) and University of Cambridge institutions and individuals: UK Home Office, implementing the Animals (Scientific Procedures) Act 1986, Amendment Regulations 2012, and represented by the local UK Home Office Inspector; the UK Animals in Science Committee; UK National Centre for Replacement, Refinement and Reduction of Animal Experiments (NC3Rs); the University of Cambridge Animal Welfare and Ethical Review Body (AWERB), the University’s Biomedical Service (UBS) Certificate Holder, Welfare Officer, Governance and Strategy Committee, Named Veterinary Surgeon (NVS) and Named Animal Care and Welfare Officer (NACWO).

### Equipment

To study the reward-effort trade-off it was necessary to develop hardware to implement different levels of effort. Approaches in the literature include lever pressing, grip force and torqued exoskeletons (Cousins et al., 1996; Pasquereau and Turner, 2013; Varazzani et al., 2015) These approaches were rejected for, respectively, the confound of delay to reward, incompatibility with binary choice designs and difficulty in training. Rather, we developed a custom effort joystick in which the resistance to movement could be computer controlled (Fig. 1A).

**Figure 1.**
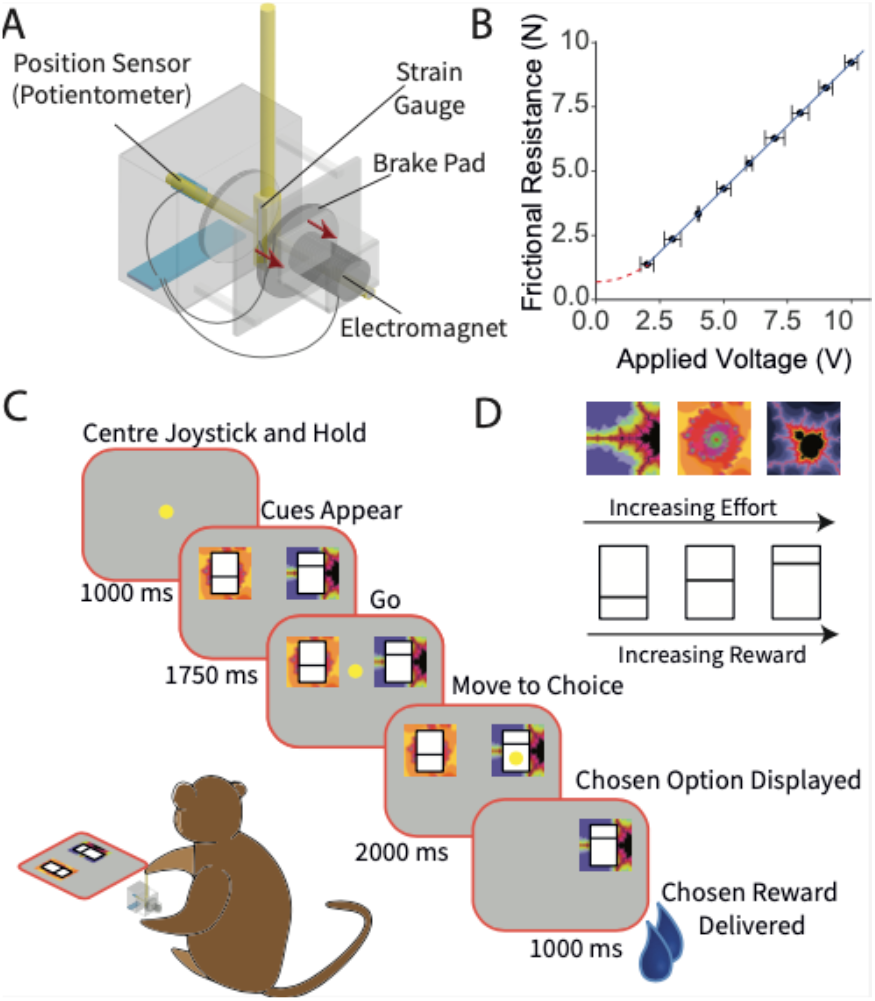
Task design. ***A***, Joystick with variable effort. ***B***, Frictional resistance as function of voltage controlling magnet strength. ***C***, Use of joystick for binary choice between options that differ in effort required. ***D***, Fractal stimuli indicating effort required (in Newton), value bars indicating reward amount (in ml).

The joystick was constructed to allow motion in a single plane of movement, from left to right, by rotation on an axis. The different effort levels were made by altering the resistance to motion by varying the strength of an electromagnet that brought the joystick into contact with a brake pad. These forces were calibrated using fixed weights under gravity: there was a linear relationship between the applied voltage and static frictional resistance (Fig. 1A). Six levels (0 – 10 V in 2V increments, corresponding to <1 N to 9.8 N; for convenience these will be considered as 0 – 10 N in 2 N steps in the analyses) of effort were used throughout the experiments. We incorporated sensors into the joystick to measure kinematic variables: capacitive touch detection, a potentiometer adapted to detect position, and paired strain gauges for detecting the directional strain on the joystick that reflects the force the monkey applied to the joystick.

For each daily experiment, each monkey was sat in its specifically adjusted primate chair (Crist Instruments) in a darkened room. A computer monitor placed 50 cm in front of the animal served to display visual stimuli. An opening on the front panel of the chair allowed the animal to express its choice by moving the joystick to pseudorandomly left-right alternating fixed positions.

The behavioral task was controlled by custom MATLAB software running on a Windows 10 computer with the Psychophysic Toolbox (Pelli, 1997) used to code the visual stimuli. Joystick position, strain and touch-state were digitized through a National Instruments Data Acquisition Card (PCI-6229) at a rate of 1 kHz. The electromagnet was similarly controlled through the data acquisition card at a rate of 250 Hz. Various amount of blackcurrant juice rewards (0 – 1 ml) were delivered by opening a solenoid (SCB262C068; ASCO) which gated a gravity-fed system.

### Binary Choice Task

The animals were trained to work for liquid rewards. The animals were mildly fluid deprived for 6 days a week during which the totality of the fluid available was provided by the performance in the behavioral task, with additional subsequent daily access to ensure the minimum daily fluid requirements were met.

The monkeys were trained to associate two-dimensional visual stimuli (Fig. 1C, D) with blackcurrant juice rewards and experienced effort over the course of >5,000 trials, including single-outcome trials. Following this training, all experimental data was gathered during a binary choice where the animal choose between two options presented simultaneously. Following option presentation, a yellow dot appeared on the screen indicating the position of the joystick and the begin of the choice phase. The animal then moved the joystick to the left or right to indicate its choice and received the reward after a 1 s delay. The chosen effort level was applied during the monkeys’ movement. Failure to make a choice resulted in the trial being repeated. Failure to complete a trial could arise for several reasons: (1) the animal did not centre the joystick in the pretrial period, (2) the animal let go of the joystick during the trial, and (3) the animal did not move the joystick to either option within a 2 s time period following the go cue. After the animal had made its choice, the stimulus of the chosen option reappeared to reinforce its association with the reward. A variable intertrial period of 1-3 s with a blank screen ensued after each trial.

### Experimental design and statistical analyses

To determine the relevant variables the monkeys used in their decision, we used a logistic model to estimate the probability of choosing a given side dependent on option variables (reward, effort) and control variables (side, previous choice location). The linear combination of these explanatory variables was related to the probability of a chosen side with the logistic function (Eq. 1) and parameters estimated within a generalized linear model.

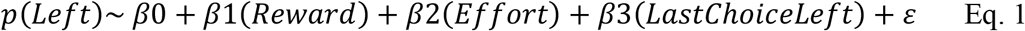

P(left):probability left, Beta0 captures side bias, Beta1 the sensitivity to reward (left minus right value), Beta2 the sensitivity to effort (left minus right), and beta3 the probability of choosing the previously chosen side, where LastChoiceLeft is a binary variable that is 1 if the left option was previously chosen, and 0 otherwise.

Further logistic regressions were used to construct a common currency between reward and effort values. Parameters were independently estimated for each day’s session.

Logistic regressions were performed using the glm (generalized linear model) function from the stats package in R, assuming an underlying binomial distribution. Relevant variables were modelled as a linear combination related to the probability of choice by the standard logistic function (y = 1/(1+*e*^-x^), with *e* being Euler’s number. The same function returned the standard error of the mean. Coefficients were standardized using the beta function in the reghelper package. Comparison of fits was done by comparing the Akaike Information Criterion (calculated by the glm.summary function), which penalizes the likelihood function by the number of terms in the model, and McFadden’s R^2^ (McFadden, 1973), a non-relative metric of fit for logistic regressions, calculated using the pR2 function of the pscl package.

Juice utility and effort disutility functions were constructed by applying random utility models on choices that differed in juice only or effort only respectively. Random utility models, or discrete choice modelling (McFadden, 1973), assume choice is dependent on the underlying utility (subjective value) of the choices and random utility (error), with decision-makers choosing the highest total (underlying plus random) utility each time. Consequently, the probability of choosing a given option is related to the difference in the underlying utility of the two options, which in the models used here they are related by the logistic function. We modelled several parametrics forms of the utility functions, including linear, power-models and quadratic, and modelled the functions non-parametrically by representing the juice/effort levels as dummy variables corresponding to the discrete levels used (0.05 ml increments for juice, 2 N for effort). In all models, daily side bias and previous side choice were included as additional explanatory variables. Parameters were estimated using the Python package BIOGEME (Bierlaire, 2020), which uses maximum likelihood estimation methods to estimate the underlying utility functions from the observed choices. To model satiety effects, we split the data of a given session into consecutive quartiles of total liquid consumption during the session; then we estimated the utility function for juice separately for each quartile. Similarly, the effort disutility function was estimated for each session quartile split by total effort exertion. In all cases, utility was normalized such that the reward utility function spanned the range 0 to 1 (by factoring out a common slope parameter).

To test whether these utility functions accurately described the animals’ choice behavior, we used the functions to predict out-of-sample choices, which compromised all options sets where both the reward and effort were different between the options.. We compared the subtractive model (reward utility minus effort disutility; u (reward) – u (cost)), hyperbolic discounting (reward multiplied by (1/(1+k*effort), where k is a scaling factor) and exponential discounting (reward multiplied by e^-k*effort^). Model comparison was by AIC and McFadden’s R^2^ (McFadden, 1973).

To determine how trial history influenced the sensitivity to effort in future trials, the logistic regression used in the first model (Eq. 1) was extended in the following terms

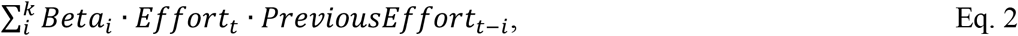

Where Effort is the effort difference on the current trial, and previous effort is the choseneffort on the ith previous trial.

To determine whether each additional previous choice term improved the fit, we compared models by computing the Akaike Information Criterion (AIC) at each step.

Given the notion that past effort experience influences an internal reference point against which current effort options are compared asymmetrically, we sought to implement a model which could identify this asymmetry. We adapted Crawford and Meng’s (2011) implementation of gain-loss utility into a random utility model as follows:

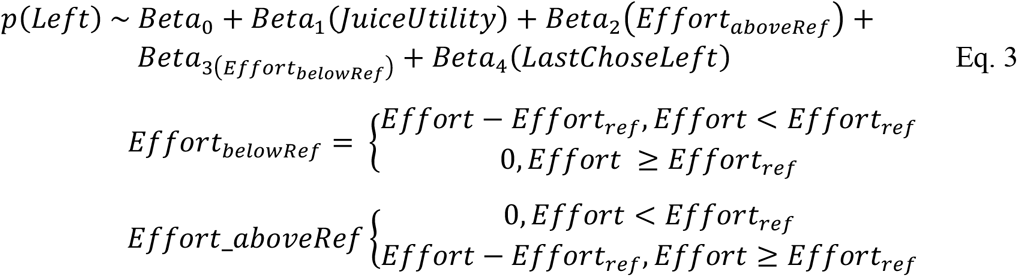

This model allows to statistically test the hypothesis that subjective effort cost is asymmetric around the reference point by testing for a difference between the standardized effort slope coefficients β2 and β3. In the first implementation of this model, effort reference was the moving average of the past 30 trials of effort.

With a view to accounting for trial-by-trial variation in the effort reference point, we developed a model of choice in which the effort reference point was learnt under a Rescorla-Wagner type reinforcement learning rule:

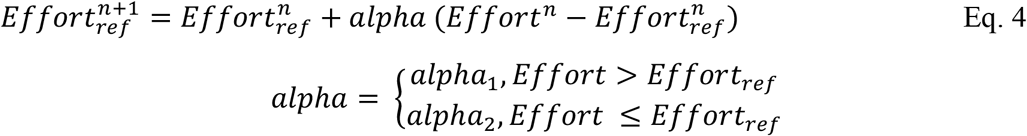

The learning rate (α) was modelled as two constants, one for efforts above reference and one for efforts below reference to allow for similar asymmetry and hypothesis testing as discussed in the estimate of the effort sensitivity parameters. Optimal learning rates were determined by grid search, with optimality defined by minimized AIC.

## Results

### Task Design

We tested choices between options that differed in the magnitude of reward the animal would obtain and the amount of effort to obtain that reward.

Following extensive training on stimuli (>5,000 trials), we presented the animals with sequences of binary choices which differed in both reward and effort levels. The choice ratio indicated how often one option was chosen over its alternative and was used to measure the relative value of the options. We were particularly interested to determine options with equal value, as indicated by their choice at equal rate (i.e. same probability of P = 0.5 of choosing each option). Through this process, we estimated and modelled the monkeys’ value of reward and effort on a single, ‘common currency’ scale.

The monkeys used the visual information to guide their decisions across all testing sessions, as demonstrated both through analysis of their choices and kinematic data. To determine the information the relevant variables used in the monkeys’ choices, we performed a logistic regression analysis on the probability of choosing the left option on the option variables (juice reward and effort) as well as other control variables (previous choice location, intercept showing left-right bias; Eq. 1).

The monkeys chose options with higher juice rewards when effort to obtain was equal (Fig. 2A, B) and chose low effort options over high effort options when juice rewards were equal (Fig. 2C, D). A positive coefficient of reward magnitude indicated that both monkeys preferred options with greater rewards (Monkey W: β=3.47, z = 160.01, p< 0.001; Monkey U: β = 1.69, z = 42.63, p < 0.001; Fig. 2E, F). The negative coefficient of effort level indicated that the monkeys preferred less effort to more effort (Monkey W: β=-0.257, z = -31.156, p < 0.001; Monkey U: β = -1.018, z = -56.6573, p <0.001). These results are consistent with rational behavior where an agent makes choices ‘as if’ maximizing subjective value by optimizing rewards (juice) at minimal cost (effort to obtain).

**Figure 2.**
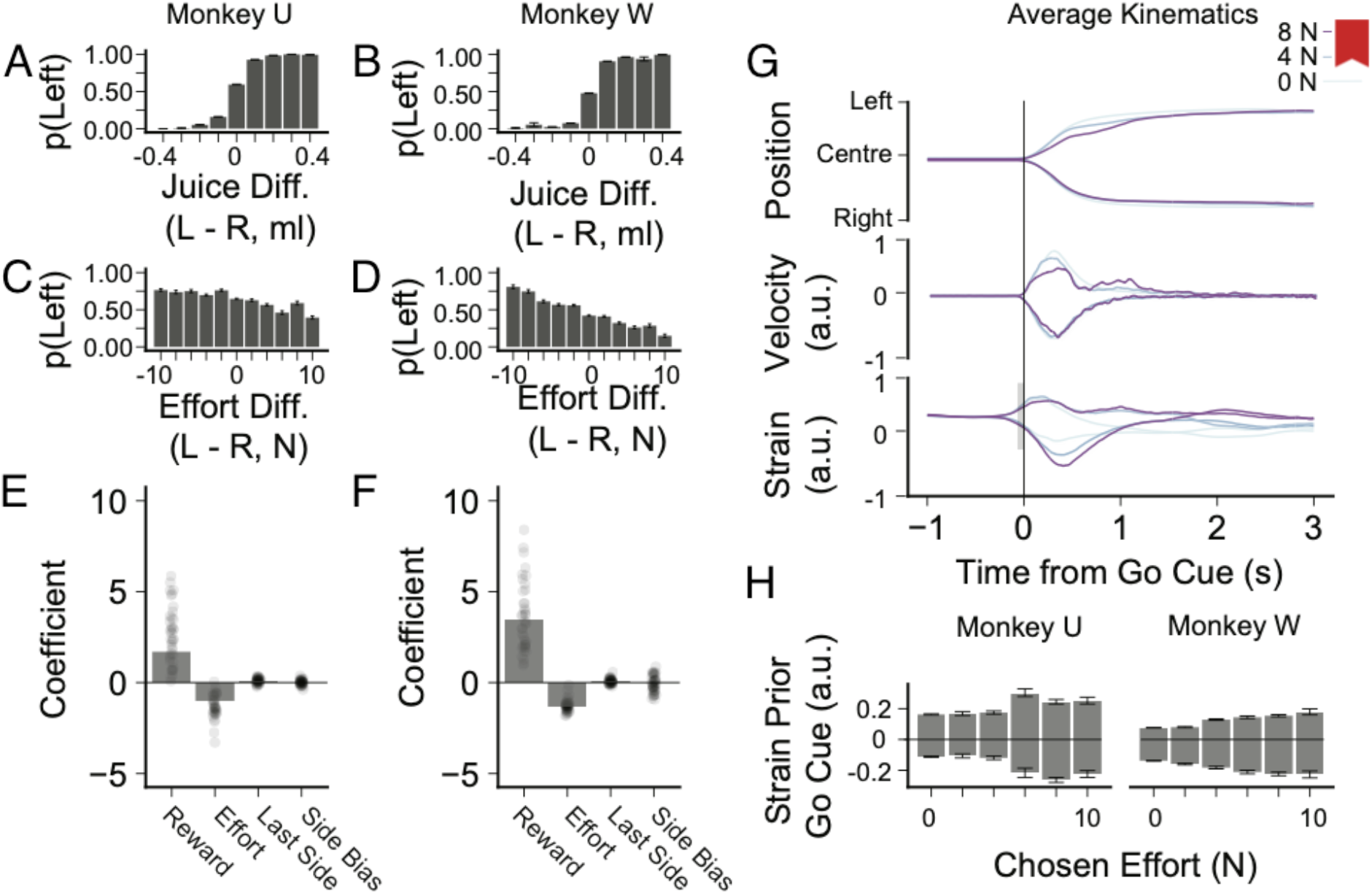
Behavioral and kinematic analysis validation and characterisation of task design. ***A, B***, Both animals chose more reward over less reward, given effort being equal. ***C, D***, Monkeys chose less effort over more effort, given equal reward. ***E, F***, Standardized coefficients from logistic regression on choosing left side, including side bias, juice and effort quantity, and bias towards previously chosen side. Column represents overall average, dots represent per-session estimates ***G***, Average profile of position, velocity and strain for Monkey U at three separate effort levels. Time zero indicates the appearance of the Go cue. ***H***, Average strain in the 100 ms before the Go cue separated by chosen side and effort level. In general, both animals were applying greater strain to the joystick for higher effort levels, reflecting preparation for the greater resistance to movement and therefore demonstrating an understanding of the effort cues. Two-way ANOVA on effort level and chosen direction; Chosen Effort Effect: Monkey U: F(5,6021) = 17.5, p < 0.001; Monkey W: F(5,4707) = 24.57, p<0.001. Error bars are SEM.

On some days there was a significant coefficient representing side bias. This left-right bias may reflect the distinct movements required for the left and right targets. Both monkeys exclusively used their right arm; therefore, a movement to the left involved wrist flexion and pronation. and elbow flexion; conversely, a movement to the right involved wrist extension and supination and elbow extension. Some forceful left movements also involved moderate shoulder adduction (right movements involved shoulder abduction) that was restricted by the walls of the primate chair. The left-right bias did not change within individual sessions, as determined by estimating the bias on a block-by-block basis and therefore in all subsequent analysis side-bias was accounted for on a day-by-day basis. Changes in side-bias may reflect the relative position of the chair to the joystick, which varied slightly day-to-day but not within sessions, with the relative position altering the balance of the movements required for left and right choices. There was a small but statistically significant bias to choose the side chosen in the previous trial (side choice hysteresis), potentially reflecting an attention bias to the previously chosen and thus rewarded side.

The monkeys’ understanding of the task was confirmed by the study of kinematic data (Fig. 2G). The joystick contained strain gauges that measured the preparatory efforts the monkeys made before the ‘Go’ cue (Fig. 2H). Averaging the directional strain in the 100 ms before the Go cue, the animals prepared to move the joystick in their chosen direction and with force that changed with the effort level of their chosen option (two-way ANOVA on effort level and chosen direction; Chosen Effort Effect: Monkey U: F(5,6021) = 17.5, p < 0.001; Monkey W: F(5,4707) = 24.57, p < 0.001). Thus, the animals seemed to be preparing to move with more force when the resistance to movement (Fig. 2H), as cued by the fractal, was higher.

### Psychometric Construction of Common Value Scale

In order to test effort cost discounting at both behavioral and neuronal levels, it is necessary to establish a common scale of value between the reward and effort. Here, we first established the viability of this method through a standard psychometric approach before extending it through random utility modelling.

To establish the equivalent value of an effort level in millilitres of juice (in the ‘common currency’ of juice), we established the indifference point (P = 0.5 choice probability) in the following way: Against a fixed magnitude of juice, at the lowest effort level, we presented a variable option (Fig. 3A). This variable option varied in juice magnitude between trials, with the same effort-level within blocks. The resulting data was fitted with a logistic function, with juice quantity as the explanatory variable. From this function, we estimated the magnitude of juice at which the variable option was chosen as frequently as the fixed option (P = 0.5 each option) (Fig. 3B). The difference in juice quantity between the indifference point and the fixed option was taken to be of the same subjective value to the monkey as the difference in the effort levels between the two options. Consistent with effort as a cost (or negative reward), the effort needed to be compensated with more juice rewards; thus effort was considered to be equivalent to negative juice quantity. To check the consistency of these measures, we repeated and measured variation across days and against different fixed options. In general, increased effort corresponded to more negative juice equivalents (Fig. 3C). The smaller monkey (Monkey W) had more negative juice equivalents for the same effort levels compared to the larger monkey (Fig. 3C left vs. right), which likely reflects the difference in their strength and thus the relative value of the effort.

**Figure 3.**
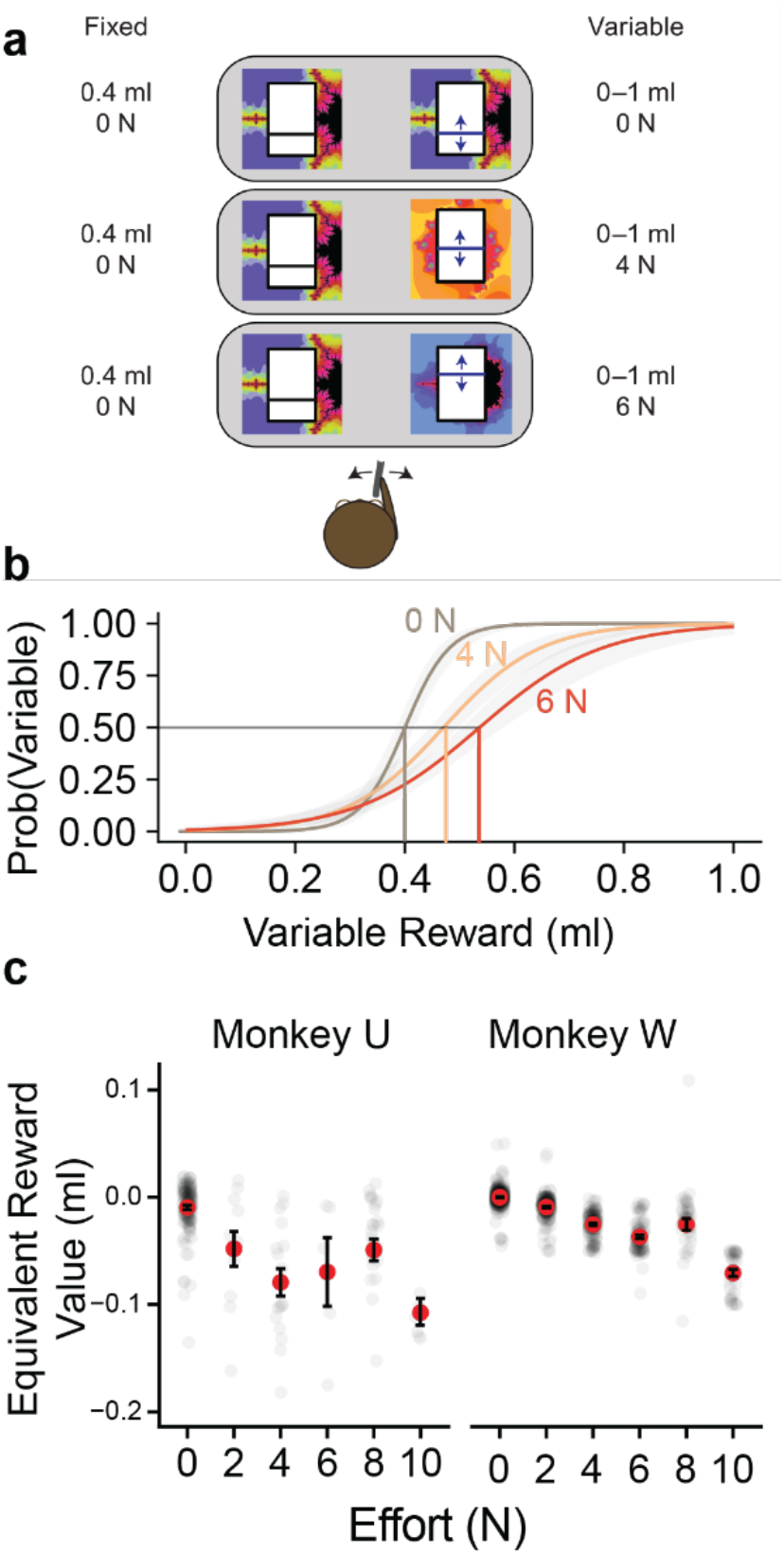
Psychometric construction of common value scale. ***A***, Task design. To establish the equivalent value of effort in units of juice the indifference point between two levels of effort was titrated. Against a fixed option (here 0.4 ml juice, lowest effort level), a variable option was presented in which the effort level and juice was changed between trials. ***B***, Example psychometric function fit for a single trial block in Monkey W (160 trials). Points and error bars represent averaged data and SEM, lines and shaded area represents fit and fit SEM. Value of effort determined from distance on juice-axis at the indifference point (choice P =0.5 of each option) for each effort level. ***C***, average equivalent juice value of each effort level for Monkey U (left) and Monkey W (right). In general, higher effort values corresponded to more negative juice values. Effort values were more negative for the smaller monkey (Monkey W), potentially reflecting a different in strength. Red dots are average over all sessions, error bars are SEM, transparent dots are session averages.

### Random Utility Modelling

Psychometric functions provide a good basis for establishing subjective value but do not completely capture subjective value. For example, if the marginal utility of juice decreases with increasing quantities of juice, the apparent cost of effort could appear lower when tested using a higher juice value as a reference. Furthermore, the apparent value of juice may shift over the course of a testing session if satiety reduces the juice value and the animal became less thirsty.

Risky-choice methods (e.g. fractile procedure; Genest et al., 2016) often reveal non-linear utility functions that may contain a mixture of concavity and convexity within the tested range. In order to obtain a more complete understanding of the subjective value of effort, it would be necessary to elicit the utility functions for gain and effort independently.

To limit the number of experimental parameters, we refrained from extensive testing of gamble options, as the additional confounds of risk preference and probability distortion might complicate the model and the understanding of the data. Therefore, we used a riskless method to establish functions for the utility of juice reward and the disutility of effort (Fig. 4A, B). In short, random utility modelling (or discrete choice modelling; McFadden, 1973) extends psychometric testing by using the choice ratio at each sampled option pair to infer the utility difference between options. In practice, we established the choice ratio between options that varied in juice but not effort, thus estimating a juice utility function. Similarly, we varied effort but not juice to estimate an effort disutility function. We used random utility modelling to fit several common utility functional forms for the reward utility and effort disutility: linear and quadratic polynomials, power function as well as non-parametrically fitting to the discrete levels of reward/effort.

**Figure 4.**
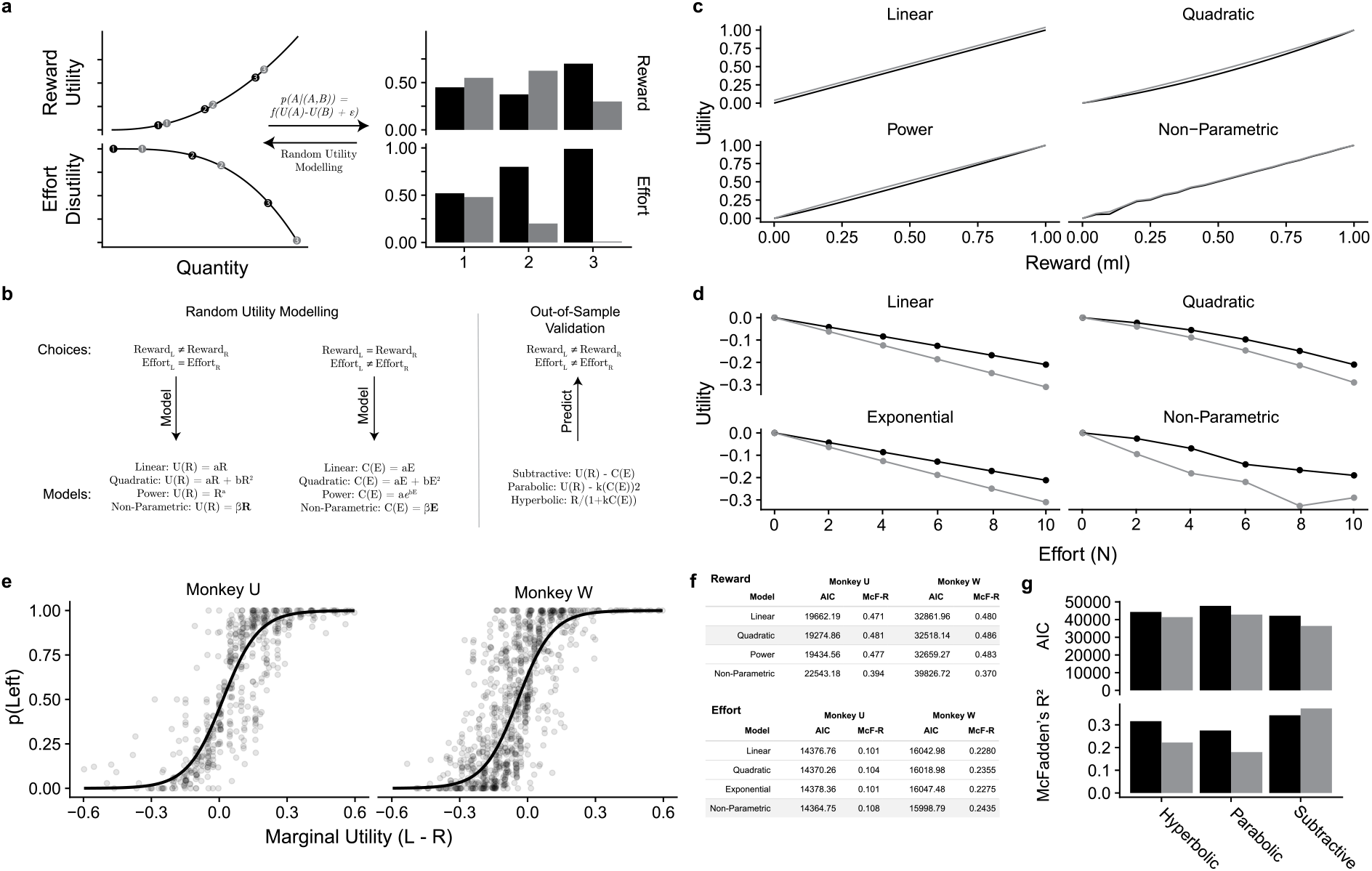
Random utility modelling and results. ***A***, Theory of Random Utility Modelling for reward and effort. ***B***, Estimation of reward utility and effort disutility functions from choice options that only differed in reward or in effort. Out-of-sample validation was conducted by predicting choices using the best fitting reward and effort disutility functions to predict choices that were not used to estimate either. ***C***, Monotonically increasing utility with reward amount; consistency across four different fitting functions. ***D***, Near-monotonically decreasing effort (dis)utility with effort (N, Newton). ***E***, Out-of-sample psychometric assessment of utility difference between leftward and rightward choices. ***F***, Quantitative comparisons of fits for the four utility functions tested. ***G***, Model fits for three commonly used combined utility models.

Through this approach, we established independent estimates of reward utility functions (Fig. 4C) and effort (dis)utility functions (Fig. 4D). For both animals and all functional forms, reward utility increased monotonically as predicted by basic economic theory. Comparing the various functional utility forms using the AIC, which penalizes model complexity, we determined that the quadratic model was the best model for both subjects. Effort disutility decreased monotonically, reflecting the choice of low effort options over high effort options. Effort disutility was best modelled non-parametrically; Monkey U was relatively insensitive to the lower efforts, whereas Monkey W was more sensitive to the lower efforts (likely owing to his smaller size), but treated the two highest effort levels similarly, as seen in the non-parametric fit (Fig. 4D). These data were replicated with all utility functions fitted (Fig. 4F).

To validate these utility estimates, we tested how well they predicted out-of-sample choices, specifically all choices that differed in both effort and reward simultaneously which therefore were not used for estimating the utility functions. We combined the reward utility and effort disutility using common methods from the literature (subtractive, hyperbolic and exponential (Białaszek et al., 2017). The subtractive model (reward utility less effort (dis)utility) provided a better fit to the empirical data than the other models we tested and a far superior fit compared to the predictive power of the juice equivalent values alone (Fig. 4G). Furthermore, the difference in utility between options predicted the choice ratio between options in a sigmoid function, confirming the model’s validity (Fig. 4E).

### Satiety Effects

The law of diminishing marginal utility captures a common-sense notion: as we have more of one thing the subjective value of an additional unit decreases. In this experiment, the advancing juice consumption of the monkeys would induce satiety and predict a decrease in the value of the juice. We inferred such a value reduction from the choice changes with the same option set between the start of the daily session and its end. To achieve this, we split each session into four consecutive quartiles by the total reward volume consumed and examined choice probabilities in each. In Fig. 5A, an example choice (0.2 ml vs 0.25 ml reward, fixed effort) is plotted. Early, when the animal is thirsty, there is a strong preference for the higher reward, but as the session continues, the animal chooses the lower option more frequently.

**Figure 5.**
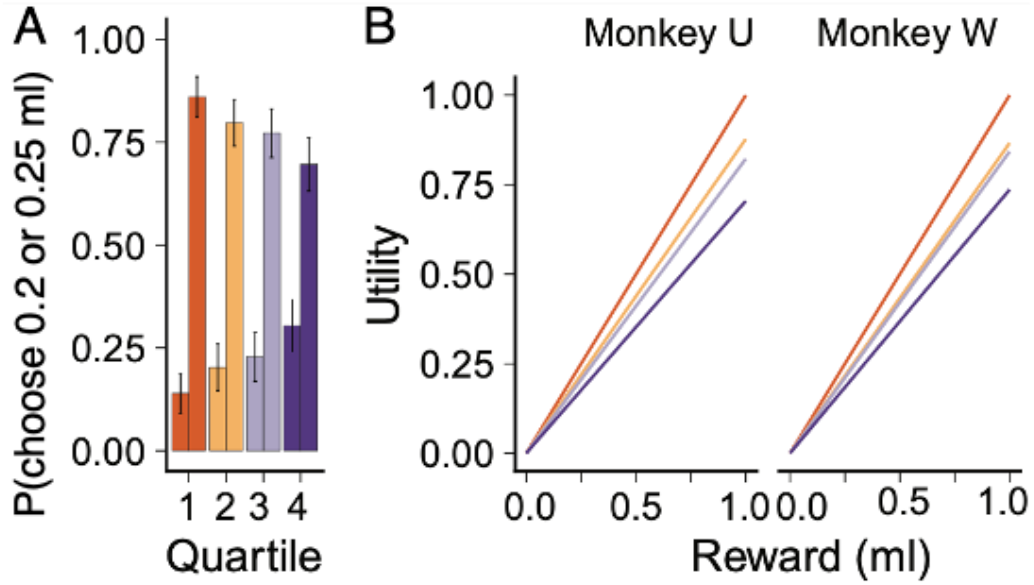
Juice consumption induces satiety effects but accumulated effort decreases. ***A***, With increasing juice consumption (quartiles 1 – 4), the choice difference between the same options narrowed, possibly reflecting satiety. For each quartile, the left column displays the probability of choosing the option yielding 0.20 ml, whereas the right column shows the choice probability for the option yielding 0.25 ml. Error bars are SEM. ***B***, Utilities for different amounts of consumed juice, as modelled by including total juice consumption as a latent variable in the juice utility model, thus quantifying the reduced juice utility as juice consumption increased. The four lines refer to the four quartiles (top to bottom).

The random utility model allowed us to quantify the change in choice ratio by estimating the reward utility function for each quartile. In both animals, the utility of reward decreased during the session (Fig. 5B), as observed in other studies (Pastor-Bernier et al., 2021).

### Adaptation to Effort

In a similar manner to satiety having an effect on the reward utility, we expected to observe a fatigue to develop over the session and increase the sensitivity to effort. Rather, we found that both subjects became less sensitive to effort over the course of each session (Fig. 6A). For example, when choosing between options requiring 0 N vs. 8 N of effort, Monkey W showed significantly stronger preference for the low effort option in earlier as compared to later daily sessions (Fig. 6A), suggesting the value difference of effort had subjectively decreased.

**Figure 6.**
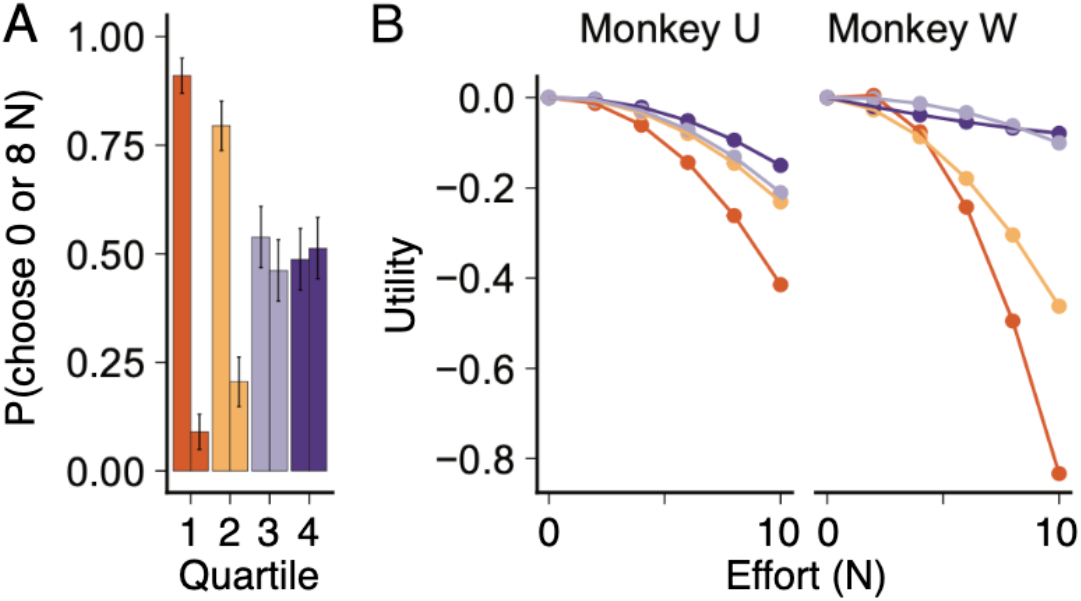
Effort choice history influences sensitivity to effort. ***A***, For each quartile of daily trials, the left column displays the probability of choosing the low effort option (0 N), whereas the right column shows the choice probability for the high effort option (8 N). ***B***, Quadratic fits of effort disutility for the four quartiles suggest effort disutility decreases with total effort expenditure over each session. The four curves refer to the four quartiles (top to bottom). Error bars show standard errors of the parameter estimate.

Splitting sessions by the cumulative effort into quartiles and using choices that only varied in effort, we estimated the effort disutility curves with a quadratic function (Fig. 6B). In both subjects, the effort disutility diminished through the session, rather than increase we would predict if the animals were fatiguing. While other studies have demonstrated that fatigue affects effort-based decision making (e.g. Hogan et al., 2020), we did not clearly observe this effect, rather the reverse.

### Effect of Previously Experienced Effort on Effort Disutility

Following this observation that sensitivity to effort was decreasing during each session, we wanted to test whether the changes in effort were dependent on the experience of previous effort, as opposed to just time in the session. To quantify the effect of previous trials, we performed a logistic regression adding the previously chosen effort level as interaction term for the variable on effort difference in Eq. 3. In a stepwise process, the explanatory power of further lagged terms was tested. Additional explanatory power was taken to be a reduction in the AIC over the previous model. This process revealed a small but significant dependence on previously chosen effort over at least the previous 27 trials in both animals.

### Asymmetric Effort Valuation relative to Effort History

A possible model to explain the diminishing effort disutility observed is the reference-dependent model. Reference dependent models pose that value is determined relative to an internal, subjective reference point. A common finding of reference models is that there is asymmetry on either side of the reference point, for example the gain-loss asymmetry in prospect theory. The evidence from the behavioral economics literature suggests that individuals are more sensitive to efforts above the reference point than to efforts of the same magnitude below the reference point (Card et al., 2012).

A simple implementation of a reference point is to use the moving mean effort of previous trials. To test this implementation, we adapted the gain-loss utility model used by Crawford and Meng (2011), with effort disutility being modelled as a piece-wise linear function, allowing for a change in slope at the reference point (Eq. 3) and using the random utility modelling techniques described above to fit the function to the data. In fitting this model to both animals, there was significant evidence of an asymmetric response to effort (Fig. 7A). For Monkey U, the parameter capturing effort sensitivity above the reference point (β, -0.44 95% CI: (−0.41, - 0.47)) was 3.6 times greater than the parameter capturing effort sensitivity below the reference point (β, -0.13; 95% CI: (−0.10, -0.16)). In Monkey W the change in slope was even greater, with the sensitivity above the reference point (β, -1.23; 95% CI: (−1.17, -1.28)) being 4.5 times the sensitivity below the reference point (β, -0.27; 95% CI: (−0.23, 0.31)).

**Figure 7.**
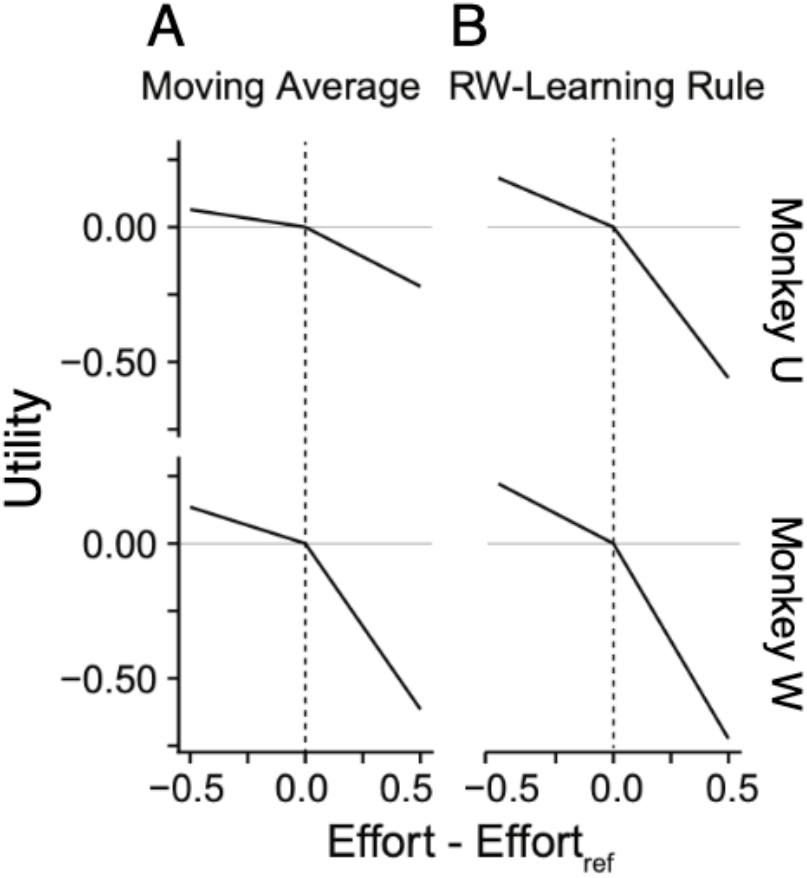
Reference-dependent effort preferences. ***A***, Effort disutility for Monkeys U and W, as scaled to the same utility as juice utility in Fig. 4. Reference was modeled as a moving average of previous chosen effort. ***B***, Reference point model from Rescorla-Wagner reinforcement learning model. In all cases, the slope above the reference point was significantly greater than below the reference point, suggesting reference-dependent preferences.

### Rescorla-Wagner Learning Model of Effort Reference Point

While the above finding is evidence for a reference-dependent effort valuation, it does not sufficiently address how the animals change their reference on a trial-by-trial basis, as the previous experiments established high effort trials have an effect on effort disutility through the session. Yet is necessary, as a basis for the neurophysiology, to understand how value changes on a trial-by-trial basis. To address this, we developed a model in which the effort-reference reflects an expectation of future evidence learned from previously experienced effort. To model learning of expected effort, we used the Rescorla-Wagner reinforcement learning model due to its inherent compatibility with temporal difference learning models, as it is based on prediction-error driven learning but has a reduced set of parameters that require estimating. In this model, the expectation of effort was updated on a trial-by-trial basis by the effort error (expected effort minus actual) multiplied by a learning rate. To allow for a difference in saliency of efforts above and below reference, analogous to the difference in saliency of losses and gains in prospect theory, the optimal learning rate was estimated separately for efforts above and below the reference point. We then used the same random utility modelling techniques with the piecewise linear function to estimate the effort disutility function. Optimal learning rates were established by a grid search method to minimize AIC.

In the optimal fits for both animals there was strong evidence that efforts above the reference point were more negatively valued than corresponding positive valued efforts the same magnitude below the reference point (Fig. 7B). The optimal learning rates also reflected that efforts above the reference point were significantly more salient than efforts below the reference point (Monkey U optimal rate above reference point: 0.089, below reference point: 0.039; Monkey W, optimal learning rate above reference point 0.29, below reference point: 0.09). Notably the ratio of learning rate above and below reference point and the slope of the utility function above and below was approximately equivalent in both animals (Monkey U: slope above reference = 2.87 × slope below reference, learning rate above reference = 2.28 × learning rate below reference; Monkey W: slope above reference = 3.26 × slope below reference, learning rate above reference = 3.22 × learning rate below reference). This suggests that the utility prediction error drives the learning of the effort reference point.

Furthermore, in this model, in which effort costs are referenced against a learned expectation of effort, better explained both monkeys’ choices versus previous models explored. It outperformed the model in which effort history was used as the reference point (Monkey U, effort history: AIC: 65770, McFadden’s R^2^: 0.445, learnt effort AIC: 65660, McFadden’s R^2^: 0.455; Monkey W effort history: AIC: 50440, McFadden’s R^2^: 0.397, learnt effort: 50070, McFadden’s R^2^: 0.401). This is potentially explained by the asymmetry of the learning rates as high efforts raise the reference effort in the learned model faster and the reference effort remained elevated even after long periods of low effort choices. This model also explained choice better in both animals versus the previous models discussed, in which juice and effort disutility were independently estimated, despite simplifying the effort disutility function to a piecewise linear function (Monkey U, disutility function: AIC: 65980 McFadden’s R^2^: 0.442, learnt effort AIC: 65660 McFadden’s R^2^: 0.455; Monkey W disutility function: AIC: 50580 McFadden’s R^2^: 0.372, learnt effort AIC: 50070, McFadden’s R^2^: 0.401).

## Discussion

In this study, we examined the cost of effort in a binary choice task and employed a proven method to estimate utility (random utility modelling) from riskless choice data. In line with numerous studies in humans, macaques and other animal models, we found evidence that effort is treated as a cost weighed against potential rewards during decision-making. Moreover, this effort cost function is not fixed, but rather dependent on an internal reference point, which is learned from previous experience.

While the psychometric method to obtain equivalent juice values for each of the effort levels was successful, the random utility model proved superior. Specifically, the subtractive model of effort discounting was the best fitting model, as predicted by economic theory (Fehr and Goette, 2007). In demonstrating that the independently estimated utility functions can be used to predict out-of-sample choices with the subtractive model, we provide direct evidence for the underlying assumption that a cost-benefit calculation underlies effort-based decision-making.

The demonstrable satiety and effort adaptation effects likely explain why the random utility model produced a monotonic decreasing effort function whereas the estimates from the psychometric analysis did not; as the monkeys consumed more juice, they were less sensitive to differences in juice quantity, reflecting diminishing utility. Similarly, increasing amounts of effort experienced diminished effort disutility. The psychometric method does not account for these changes and therefore efforts tested towards the end of the session would be expected to have smaller equivalent juice values than ones tested earlier in the session, adding significantly to the variance. Likewise, the value of reward is not constant over the course of a given session.

Returning to the example of the introduction, walking to the kitchen for breakfast and hiking up a mountain on holiday involve wildly different amounts of effort expenditure yet the brain is able to decide to undertake these actions implying flexible decision mechanisms. Expectation is likely a major component of this flexibility: in economics, the expectation of reward and cost has been demonstrated to influence choice behavior (Kahneman and Tversky, 1979). The most successful explanation of effort disutility we found to explain the animals’ choices was the reference dependent model in which the reference is an expectation of effort learned from previous trials. The dependence on expectations has been demonstrated in humans by comparing unexpected and expected changes in wage on work behavior in Swiss bicycle messenger service in Zurich (Fehr and Goette, 2007). In biology, adaptive patterns in neuronal codes are common, as range-adaptation is an inherent feature of the brain. Sensory neurons tune their firing to match the distribution of sensory signals, and recent evidence suggests similar patterns exist in neurons encoding value, including midbrain dopamine neurons (Tobler et al., 2005; Kobayashi et al., 2010; Louie et al., 2015; Burke et al., 2016). Behaviorally, many psychological experiments have demonstrated that effort generalizes and adapts across different behaviors like the adaptation to effort observed here (Eisenberger et al., 1979, 1989).

One possibility not accounted for here is that low effort options, particularly the lowest effort option, may have positive utility. If the average effort required to obtain a reward is high, a low effort option may be perceived as a rest from effort and therefore may have increasing or even positive utility that is not revealed in these analyses due to anchoring the zero-effort option to zero utility. It is difficult to assess this effect with this task as the animals may also rest by skipping trials and it is not possible to correlate this effect with fatigue as the effort-adaptation effects appear to obscure any fatigue effects. It was necessary to define an anchor point on which to scale utility as utility is only unique up to positive affine transformations. Therefore, the positive or negative value of utility is only in relation to this anchor point.

In the context of this task design, the monkeys can only learn about what effort to expect from their own experience of the effort and the presentation of previous cues. In these experiments and models, we investigated the role that the order of presentation of effort cues and the experience of effort has on the subjective value of effort. In line with the economic literature on human behavior, we found monkeys’ effort preferences were reference-dependent and shifted following changes to the range of efforts experienced in the task. As the monkeys experienced effort, their choices became progressively less sensitive to the effort difference between options. Modelling this as a prospect-theory like reference-dependent function suggests that the monkeys were particularly sensitive to efforts above their expected effort and relatively insensitive to efforts below their expected effort.

The learning model of effort, in which the reference point is an expectation of effort learnt from previous trials driven by a prediction-error mechanism, improved over models in which effort was a fixed-cost throughout the session. This model, which is an adapted Rescorla-Wagner type learning rule, is a model of reinforcement learning. Here the similar ratio of the learning rates above and below the reference point, with the relative utilities of effort above and below the reference point suggests that effort utility errors drive the learning of expected effort. Similar reinforcement learning models have been a very successful framework for understanding neuronal activity within brain areas involved in decision-making, including the midbrain dopamine neurons. Midbrain dopamine neurons encode reward prediction errors and optogenetic experiments, in which dopamine neuron activity was manipulated to enhance or reduce prediction errors, observe changes in reward learning behavior that demonstrate a causal link between dopamine prediction errors and reward valuation (Stauffer et al., 2016). We did not exhaustively model all possibilities of adaptive functions, only showing that the learning rule, which accounts for potential asymmetry outperforms a rolling-average: we did not consider any model-based learning paradigms, nor were we able to examine inter-session effects.

A learned effort reference point is evolutionary useful: theory and modelling suggest that an appropriate reference point that reflects realistic effort requirements supports optimal effort-based decision-making (Rigoli and Pezzulo, 2022). Unlike in this task, expectations may be learned from more than previous experience alone; the human literature provides good evidence that effort expectations learned from social information influence effort-based decisions (Card et al., 2012).

In neuroscience, computational modelling, notably of activity in the anterior cingulate cortex, an area known to be involved in effort valuation, have directly addressed adaptive effort effects (Verguts et al., 2015). These models suggest that effort is both learnable and context-dependent and therefore expectations of future effort influence the value of effort, potentially suggesting the anterior cingulate cortex is a neural substrate for effort expectation calculations. The anterior cingulate cortex is involved in decision-making circuitry, for example it exerts influence over dopamine neurons in the ventral tegmental area via monosynaptic glutamatergic connections, suggesting such adaptive effort values may be represented in dopamine neurons as well (Beier et al., 2015).

Reference dependent efforts may explain why previous studies investigating the link between phasic dopamine and effort costs have not found consistent coding of effort costs following presentation of effort-predicting cues (Gan et al., 2010; Pasquereau and Turner, 2013; Varazzani et al., 2015). If the subjective value of effort is reference-dependent in the manner suggested here, then pseudorandom ordering of effort levels will bias the reference effort level towards the high end of the range of efforts, due to the asymmetry in value driving the learning of expected effort. When efforts are compared to a high effort reference point, most of the effort range will lie below the reference point, in which the relative value between effort levels is diminished and therefore little-to-no encoding of effort found in phasic dopamine. This is supported by the finding of Gan et al. (2010), in which there were sudden changes in the effort level were accompanied by marked modulation of dopamine release in the nucleus accumbens, suggesting dopamine may encode value relative to the expected value of effort.

Overall, these experiments suggest that monkeys’ subjective value of effort is reference-dependent, and that the reference point is, at least in part, an expectation of the average effort level formed from experience in previous trials. Such effects are necessary to control and understand when probing the relationship between midbrain dopamine neurons and effort cost encoding.

## Acknowledgements

We thank Aled David and Christina Thompson for animal and technical support and Philipe Bujold and Fabian Grabenhorst for discussions on the design and statistics of this experiment. This study was supported by Wellcome Trust (WT 095495, WT 204811, WT 206207), European Research Council (ERC; 293549) and US National Institutes of Mental Health (NIMH) Caltech Conte Center (P50MH094258). For the purpose of Open Access, the authors have applied a CC BY public copyright licence to any Author Accepted Manuscript version arising from this submission.

